# Perfusion MRI using endogenous deoxyhemoglobin as a contrast agent: preliminary data

**DOI:** 10.1101/2020.08.21.255802

**Authors:** J. Poublanc, O. Sobczyk, R. Shafi, J. Duffin, K. Uludag, J. Wood, C. Vu, R. Dharmakumar, J.A. Fisher, D.J. Mikulis

## Abstract

**Background:** The paramagnetic properties of deoxyhemoglobin shorten T2* as do gadolinium based contrast agents. Induction of abrupt changes in arterial deoxyhemoglobin concentration ([dOHb]) can simulate intra-vascular injections of gadolinium for perfusion imaging.

**Aim:** To demonstrate the feasibility of making rapid changes in pulmonary venous hemoglobin saturation and employing the resulting changes in T2* to calculate flow metrics in the brain.

**Methods:** A gas blender with a sequential gas delivery breathing circuit and software enabling prospective arterial blood gas targeting was used to implement rapid isocapnic lung changes in the partial pressure of blood oxygen (PaO_2_). Lung PO_2_ was initially lowered to induce a low baseline [dOHb]. PaO2 was then rapidly raised to PaO_2_ ∼ 100 mmHg for 10 seconds and then rapidly returned to baseline. Blood oxygenation level dependent (BOLD) MRI signal changes were measured over time.

**Results:** Arrival delay, signal amplitude and change in BOLD discriminated between large arteries, tissue and veins. The median half-time of BOLD signal in the middle cerebral artery was 1.7 s, indicating minimal dispersion confirming effective rapid modulation of pulmonary venous PO2. The contrast-to-noise ratio in the cortex was 3. Calculations of arrival delay, cerebral blood volume, mean transit time and cerebral blood flow were within normal ranges from published literature values.

**Conclusion:** Non-invasive induction of abrupt changes in [OHb] may function as a novel non-invasive vascular contrast agent for use in perfusion imaging.

## Introduction

Blood Oxygenation Level-Dependent (BOLD) sequences are sensitive to distortions in the static magnetic field caused by the concentration of paramagnetic moieties such as deoxyhemoglobin ([dOHb]) and gadolinium-based contrast agents^1^. The time constant of the exponential decay of the BOLD signal T2* is inversely proportional to the concentration of the paramagnetic moieties. There are near linear > 3-fold decreases in intra-vascular T2* with increasing [dOHb] to 50% ^2^,^3^ at 3 Tesla ^4^--half the effect of the BOLD signal lowering effect of GBCAs. Thus [dOHb] would have the advantage of being an intrinsically generated molecule with the potential of a formidable contrast agent.^5^ The [dOHb] would be generated by changing the PO_2_ in the lungs. The range of PaO_2_ affecting the [dOHb] is in the range from ∼100 mmHg (normoxia, with absent [dOHb]) to about ∼30 mmHg (50% [dOHb]) ^6^; notably excluding hyperoxic levels of PO_2_ > 100 mmHg, where arterial [dOHb] is normally absent.

The generation of dOHb as a contrast agent has seldom been exploited for hemodynamic studies due to the difficulty in precisely targeting arterial PO_2_ (PaO_2_) by the inspired PO_2_. This difficulty is compounded by the requirement of simultaneously and strictly maintaining a constant arterial PCO_2_ (PaCO_2_) in order not to disturb the cerebral blood flow. Once the means of achieving these targeted lung gas transitions is solved, the physiology of gas-blood equilibration is very advantageous to the control of contrast formation. Inspired gas instantaneously distributes widely throughout the lungs, roughly matching pulmonary blood flow. The very large diffusion interface (50-75 m^2^, about half the size of a tennis court) then results in a near instantaneous equilibration of the lung PO_2_ with that of mixed venous blood, and hence the corresponding [dOHb] which is conducted directly to the systemic arterial circulation.

For over 20 years rapid precisely targeted lung and arterial blood PCO_2_ have been induced by the method of sequential gas delivery (SGD) ^7^. Our aim was to use the SGD to implement abrupt, targeted transitions of PO_2_ in the lung, (while maintaining PCO_2_ constant) to generate target [dOHb] as a MRI contrast and thus enable new measures of cerebral hemodynamics. For this proof-of-concept protocol, we elected to establish a hypoxic baseline, PO2 < 100 mmHg with the contrast generated by a transient increase in PaO_2_ and drop in [dOHb].

Between December 18, 2019 and January 28, 2020 four of the Toronto-based investigators were scanned during the MRI protocol development sessions. Soon thereafter the hospital closed all research studies to focus on COVID-19 related patient care. Despite the small variations in protocols, the investigators found the generated data remarkably consistent, and of sufficiently high contrast to noise ratio (CNR) to merit an early alert to the imaging community. In this communication, we are able to report: 1) the successful implementation of an abrupt 10 s reduction in [dOHb] and its BOLD response; 2) the BOLD contrast-to-noise ratio (CNR) resulting from the change in [dOHb]; 3) the degree of [dOHb] bolus dispersion upon arrival in arteries of the brain constituting an effective arterial input function; 4) calculation of standard blood flow parameters that are consistent with those in previous publications.

## Methods

As these methods for inducing [dOHb] changes are totally unprecedented, we were obliged to perform initial exploratory studies to develop the study protocols prior to the IRB application. IRB considered these studies developmental nature and that all subjects were investigators and thus well informed, and did not require full IRB review. All scanned investigators were in good health without contraindications to MRI environment and brief episodes of mild hypoxia. Each provided signed consent for the studies.

### Respiratory protocol

An investigational computerized gas blender, the RespirAct™ RA-MR (Thornhill Medical, Toronto, Canada), was used to prospectively target lung PO2 while maintaining normocapnia independently of ventilation and target lung PO2.^8,9^ End-tidal PO2 (PETO2) was lowered to a baseline of about 55 mmHg (hypoxia baseline) while maintaining normocapnia in order to minimize the changes in arterial PCO2 (and hence its effect on cerebral blood flow). Hypoxia was maintained for 60 to 90s followed by a gas challenge with the PETO2 target set to 100 mmHg for 10s to transiently saturate hemoglobin with O2 before returning to baseline. This normoxic challenge and return to hypoxic baseline was then repeated 3 to 4 times during each trial (Figure 1).

**Figure 1:**
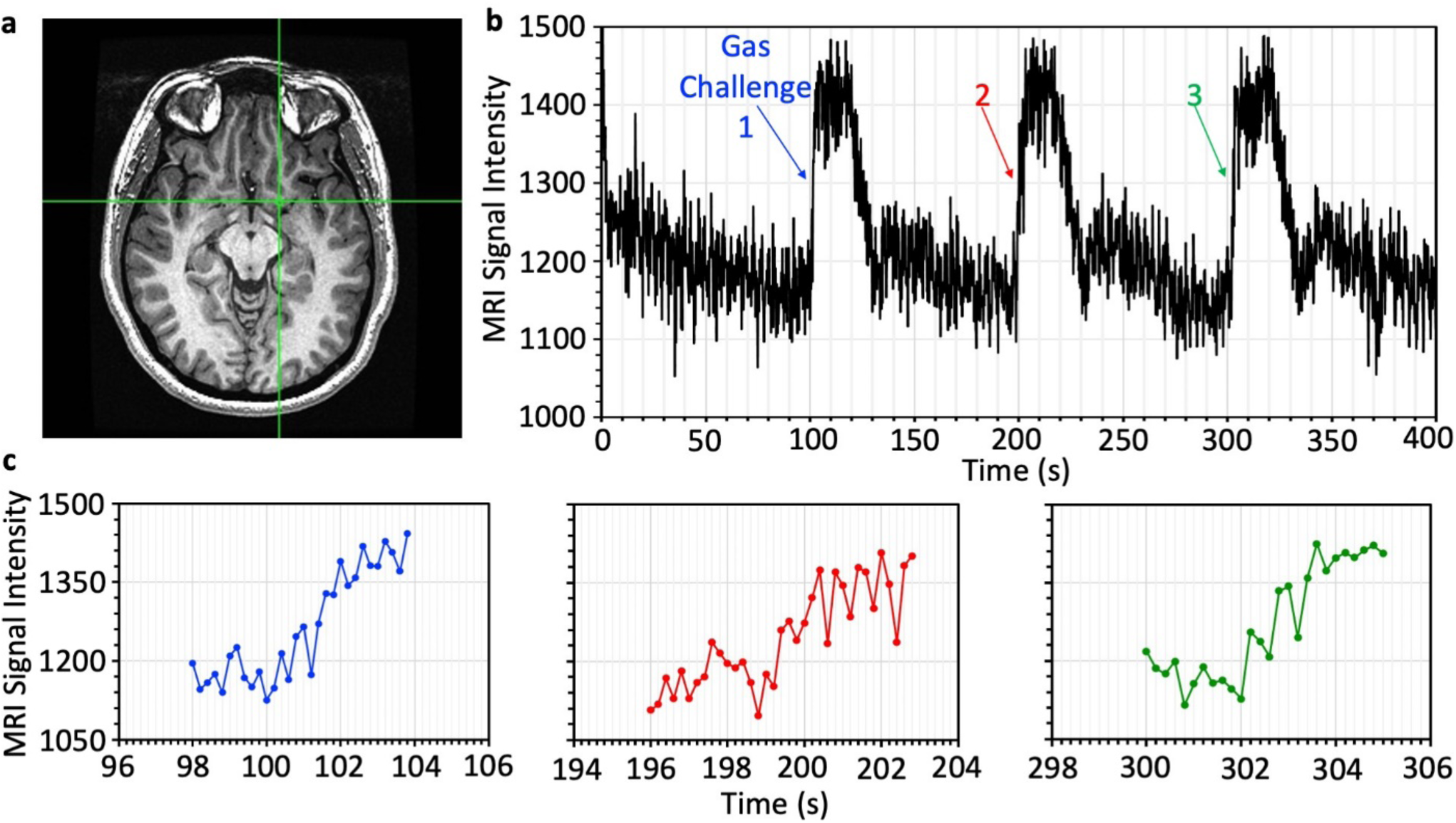
Gas challenge and BOLD arterial signal. The protocol was first carried out with a short TR = 200 ms for measuring response time. A voxel overlying the middle cerebral artery (green cross–hairs) with a large change in BOLD signal was chosen to represent the arterial signal rise time. The first 3 normoxic challenges are shown. a: Location of the voxel. b: BOLD signal vs. time for 3 consecutive challenges. c: Expanded time scale of the BOLD signal change in B from baseline to normoxia. Fitting a first order exponential to the rise in the BOLD signal results in time constants of 1.21 s, 1.67 s and 2.10 s respectively.

### Magnetic Resonance Imaging

BOLD images were acquired on a 3T MRI system (Signa HDx - GE Healthcare, Milwaukee) with interleaved echo-planar acquisition during PETO2 manipulation. The first experiment used a short TR of 200ms, 3 mm isotropic voxels with 3 contiguous slices positioned over the middle cerebral arteries (Figure 1). We identified voxels with large CNR and fit a first order exponential to the rise in BOLD and derived a time constant as a measure of dispersion. A second experiment with 4 normoxic transitions, consisted of 37 contiguous slices with an isotropic resolution of 3mm, field of view of 19.6cm, and TR/TE 2000/30 ms. A high-resolution T1-weighted SPGR sequence was acquired for co-registering the BOLD images and localizing the arterial and venous components. SPGR parameters included: 176 slices of 1mm thick partitions, in-plane voxel size of 0.85×0.85mm, field of view of 22cm, and TR/TE = 7.88/3.06 msec.

### Conversion of end-tidal partial pressure of O2 (PETO2) to [dOHb]

The recorded end-tidal PETO_2_ was converted to SaO2 using the Hill equation normal sea level physiologic parameters ^6^:

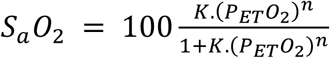

*Where n* = −4.4921 . *pH* + 36.365; *K* = 5.10^−142^.*pH*^157.31^; *pH = 7*.*4*

[dOHb] = [OHb] x (1-SaO_2_)

### Image Analysis

All image analysis was performed using Matlab 2015a and AFNI ^10^. The 3-slice dataset with a short TR of 200ms was visually inspected and one voxel with high CNR localized over the middle cerebral artery was chosen to analyze the rise-time (*τ*) of the BOLD signal. *τ* is the time constant of the exponential function fitted to the BOLD signal. The pre-processing steps for the whole brain dataset with TR of 2000ms included slice time correction, volume re-registration, and fourth degree polynomial detrending. SaO_2_ was resampled and interpolated to TR intervals and time-aligned to the whole brain average BOLD signal.

The baseline BOLD signal (S_0_) was then defined as the mean of the BOLD signal (S_t_) over five time-frames including the duration of the baseline before the first bolus, 10 sec before the second, third and fourth boluses, and 10 sec before the end of the protocol. The scaled BOLD signal (S_c,t_) could then be calculated as 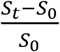.

### ΔBOLD, CNR and TD

A time delay (TD) map was calculated using cross-correlation between *S*_*c,t*_ and multiple *S*_*a*_*O*_2_ curves time shifted from 0 to 5 sec by intervals of 0.1 s. The time shift needed to obtain maximum correlation (R) with *S*_*c,t*_ was extracted for each voxel to generate a time delay.

*S*_*c,t*_ was regressed against the voxel-wise shifted 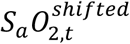 to calculate the slope of regression:

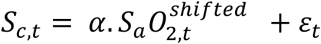

with *α* = slope of the regression and *ε*_*t*_ the residuals.

Using the slope of the regression *α*, BOLD signal change (ΔBOLD) and CNR were computed as:

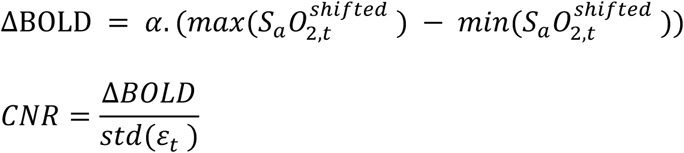

In order to examine whether signal averaging would improve bolus definition, CNR was calculated for 1, 2, 3 and finally all 4 gas challenges: the range of each segment was defined as 0-190 sec, 0-290 sec, 0-390 sec, 0-490 sec.

### Tracking of contrast in arteries and veins

Arterial and venous voxels were extracted using Δ*BOLD*, R, and TD. Arterial voxels were defined as all voxels with Δ*BOLD* > 20%, R > 0.8 and 0 < TD < 1.5 sec. Venous voxels were defined as all voxels with Δ*BOLD* > 20%, R > 0.8 and 3 < TD < 5. BOLD time series was averaged within the arterial and venous components over the four boluses.

The SaO2 was also averaged over the four boluses.

Cerebral blood volume (CBV)

Quantitative CBV was calculated as follows:

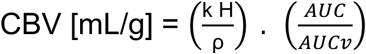

Tissue density: ρ = 1.04 g/cc

Difference in hematocrit in large vessels and capillaries: kH = 0.73 kH/ρ = 0.705 cc/g

AUC is the area under the curve of the BOLD signal (S_t_) over the entire protocol. AUCv is the area under the curve for a voxel containing blood only with large AUC. We set this value to correspond to the 5th percentile of the largest AUC.

### Mean transit time (MTT) and cerebral blood flow (CBF)

Standard tracer kinetic modeling was used to calculate MTT and CBF as per:

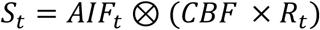

Subscript t indicates that the variable is a function of time.

S_a_O_2_ was used as the arterial input function (AIF_t_) and was scaled to have an AUC equal AUCv.

*R*_*t*_ = *e*^−*t*/*MTT*^ was used for the residue function. It is equal to 1 at time 0 and set to 0 at time equal to 5 x MTT. CBF and MTT can then be determined using least square fitting procedure. MTT was bound between 0 and 7s.

### Gray and white matter measurements

The T1-weighted images were segmented into gray matter, white matter and cerebrospinal fluid using spm8 software package ^11^. The probability density maps obtained were thresholded at 0.8 to generate a mask for gray matter (GM) and white matter (WM). One layer of peripheral voxels was eroded from the white matter mask to minimize partial volume effects.

Gray and white matter mask were used to calculate average values of BOLD signal, CNR, TD, MTT, CBV and CBF. A paired t-test was also computed to assess the change in CNR for each set of merged boluses.

## Results

Figure 1 illustrates the extent of dispersion inclusive of that in the lung, confluence in the pulmonary vein, wash-in and wash-out of the left ventricle, and any mixing in the aorta and extracranial arteries. There is also remarkable consistency in the successive gas challenges and resulting BOLD changes with little drift (Figure 1 b). Figure 2b illustrates that the distribution of early arriving large amplitude voxels (red) are consistent with the locations of middle cerebral arteries, and the later arriving large amplitude changes, colored blue, are consistent with the location of the major veins and venous sinuses. The CNR was sufficiently large that averaging results of gas challenges resulted in minimal improvement of CNR. Although changes in CNR for each additional bolus are not large, a Bonferroni corrected paired t-test showed that all of the changes were significant (p < 0.001) for both gray and white matter.

**Figure 2:**
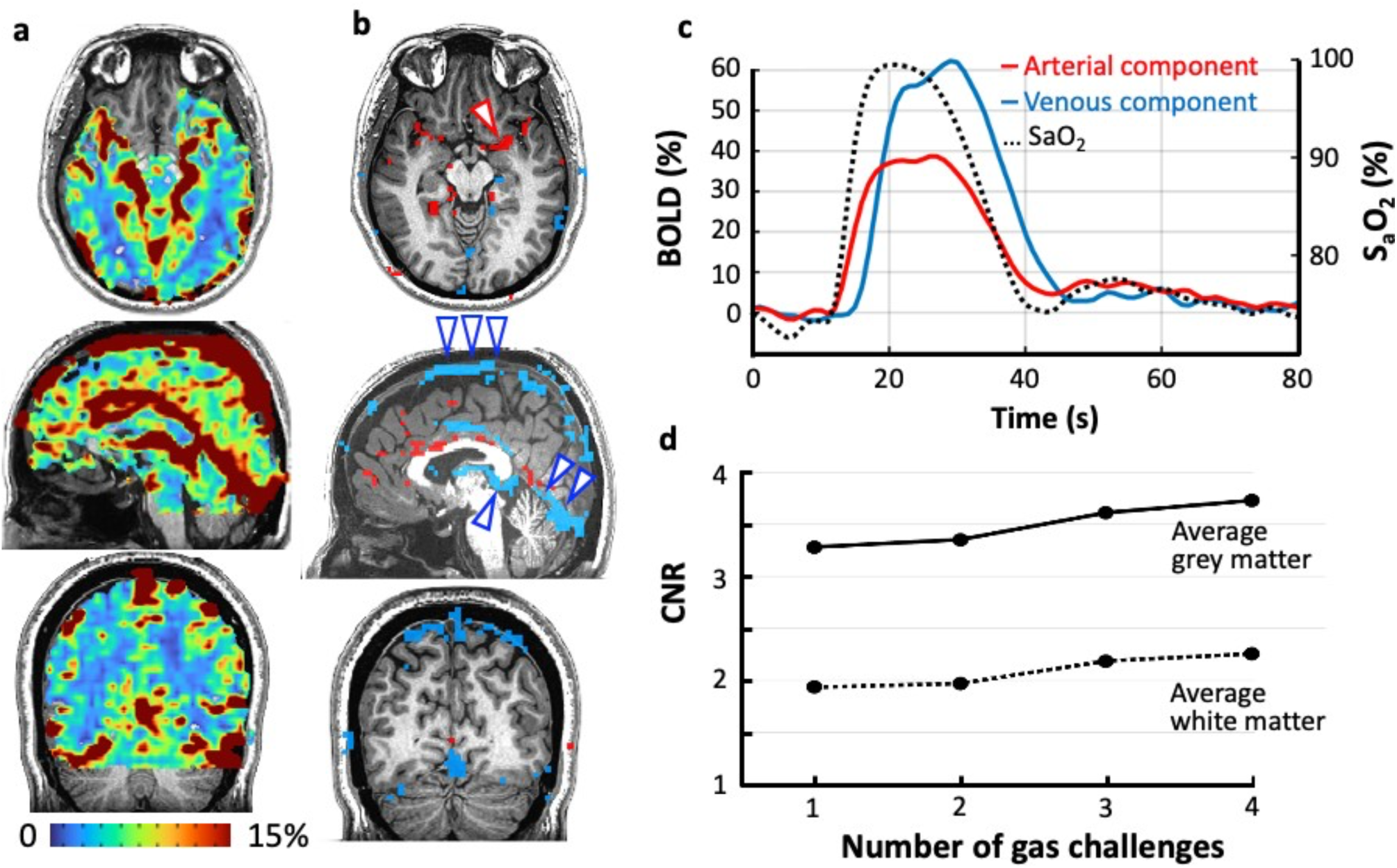
Changes in BOLD signal due to changes in deoxyhemoglobin concentration [dOHb]. The protocol was carried out with TR = 2000 ms. **a**: Maps of BOLD signal change (Δ BOLD). **b**: Large arteries and veins can be distinguished based on arrival time differences (see c) and amplitude of the paramagnetic bolus. Identification of large veins is confirmed on visual inspection of the maps as voxels that contain the highest volume of blood and least volume averaging with adjacent low blood volume tissues will have the largest signal change. Late filling voxels co-localized with known large sinovenous structures: superior sagittal sinus (3 blue wedges), straight sinus (2 blue wedges), and Great vein of Galen (one blue wedge). **c**: Graph showing bolus arrival times. The SaO2 curve was calculated from (PETO_2_) data synchronized with the rise in arterial BOLD signal. **d**: The contrast to noise ratio (CNR) for 1 gas challenge, as well as the CNR after averaging over 2, 3 and 4 gas challenges for gray matter and white matter. Averaging over successive challenges adds little to CNR.

Figure 2a shows BOLD signal change clearly distinguished large vessels, gray matter, and white matter. Figure 2c shows the average BOLD signal over arterial and venous voxels separately. This average was calculated over the four gas challenges. Delay maps show close matching to tissue types (Figure 3). Maps displaying CBV, MTT and CBF calculated from the generated data are shown in Figure 4. Values of perfusion metrics are listed in table 1 (under dOHb bolus) showing good agreement with published literature values.

**Table 1.**
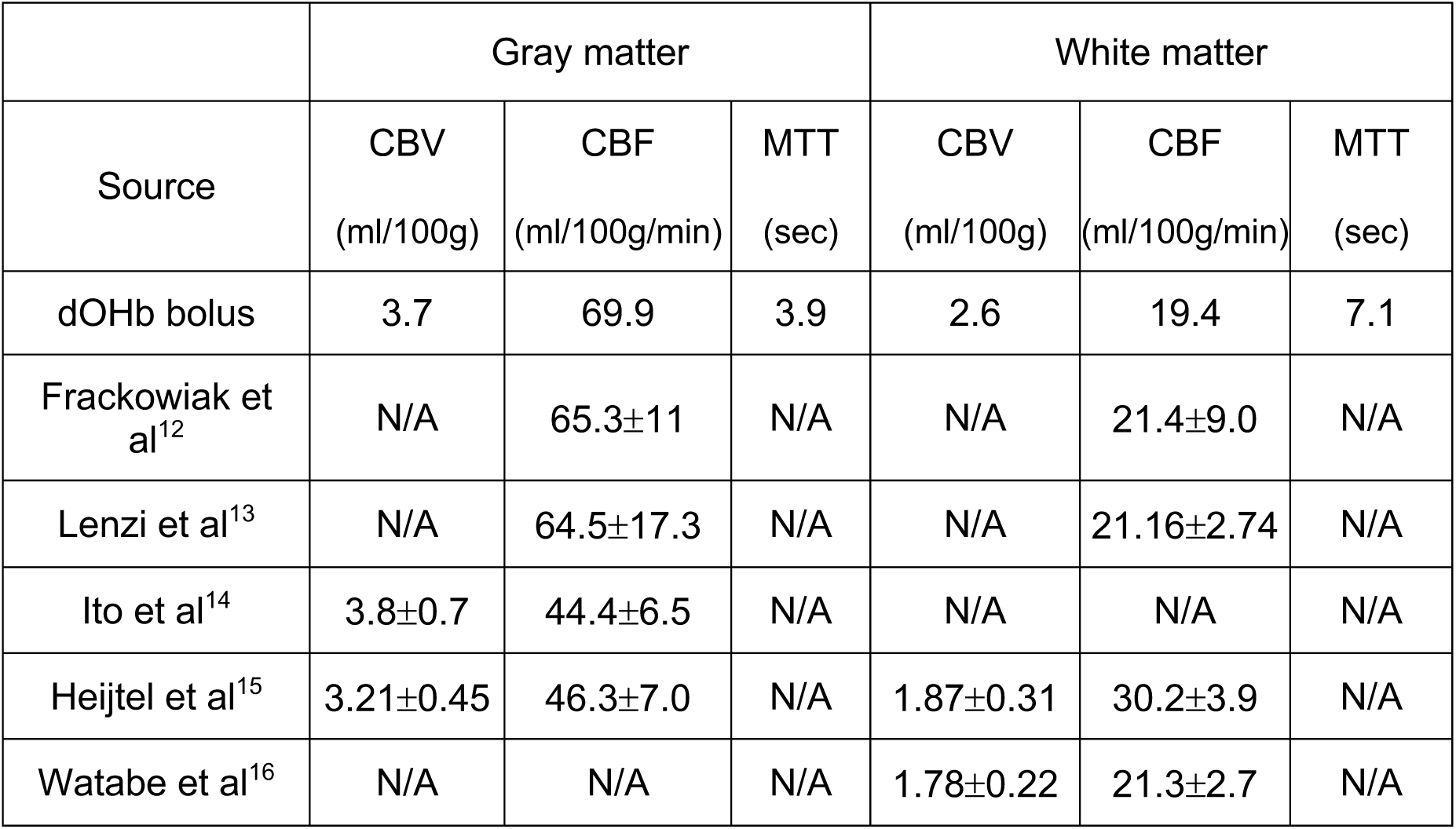

|  | Gray matter |  |  | White matter |  |  |
| --- | --- | --- | --- | --- | --- | --- |
| Source | CBV<br>(ml/100g) | CBF<br>(ml/100g/min) | MTT<br>(sec) | CBV<br>(ml/100g) | CBF<br>(ml/100g/min) | MTT<br>(sec) |
| dOHb bolus | 3.7 | 69.9 | 3.9 | 2.6 | 19.4 | 7.1 |
| Frackowiak et al<sup>12</sup> | N/A | 65.3±11 | N/A | N/A | 21.4±9.0 | N/A |
| Lenzi et al<sup>13</sup> | N/A | 64.5±17.3 | N/A | N/A | 21.16±2.74 | N/A |
| Ito et al<sup>14</sup> | 3.8±0.7 | 44.4±6.5 | N/A | N/A | N/A | N/A |
| Heijtel et al<sup>15</sup> | 3.21±0.45 | 46.3±7.0 | N/A | 1.87±0.31 | 30.2±3.9 | N/A |
| Watabe et al<sup>16</sup> | N/A | N/A | N/A | 1.78±0.22 | 21.3±2.7 | N/A |

**Figure 3:**
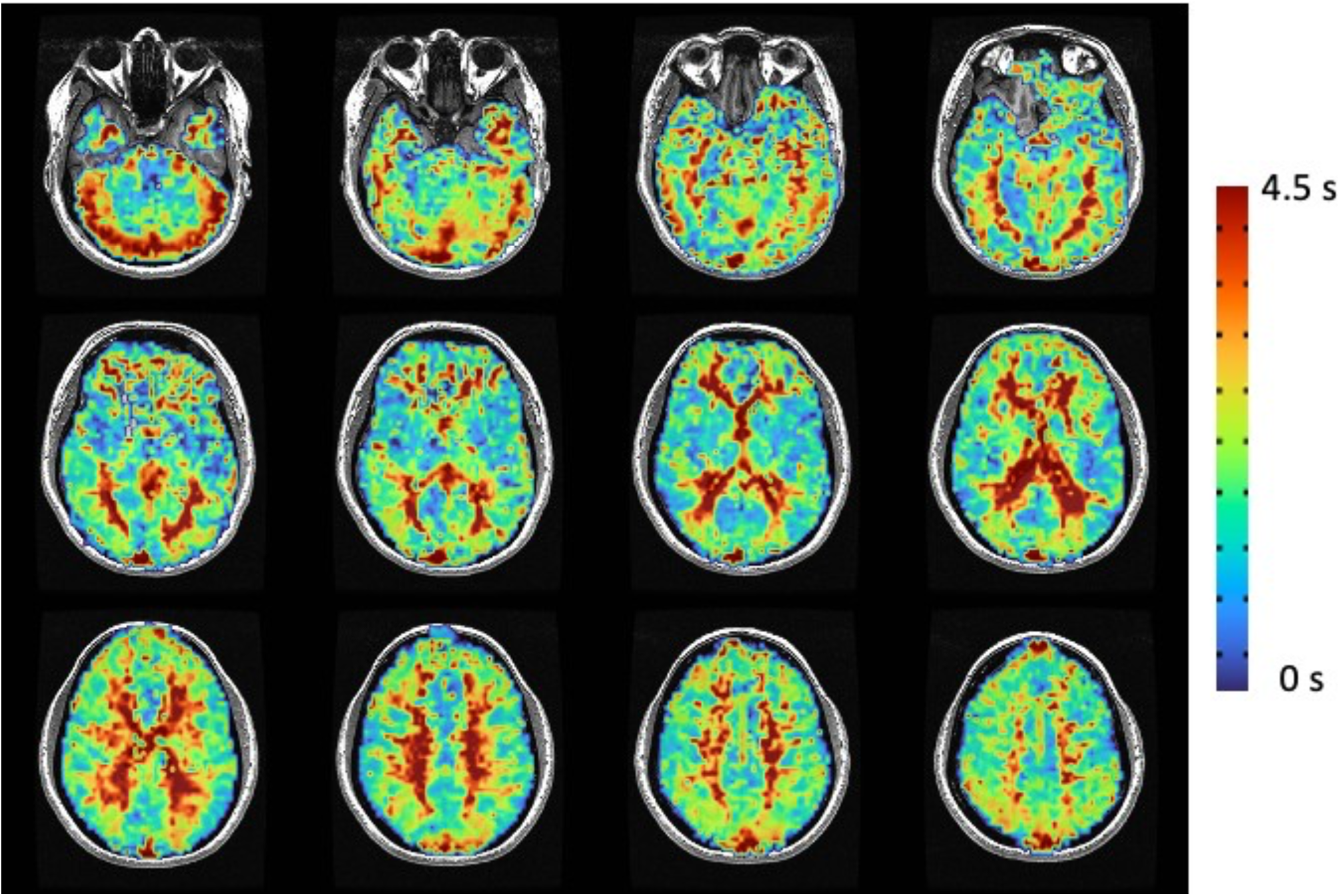
Time Delay (TD) map The map shows sharp delineation between initial arrival of the oxyhemoglobin bolus in gray matter (blue/green), followed by subcortical and deep white matter (green/yellow), and late arrival in periventricular white matter and veins (orange/red) consistent with the known cerebral vascular kinetics.

**Figure 4:**
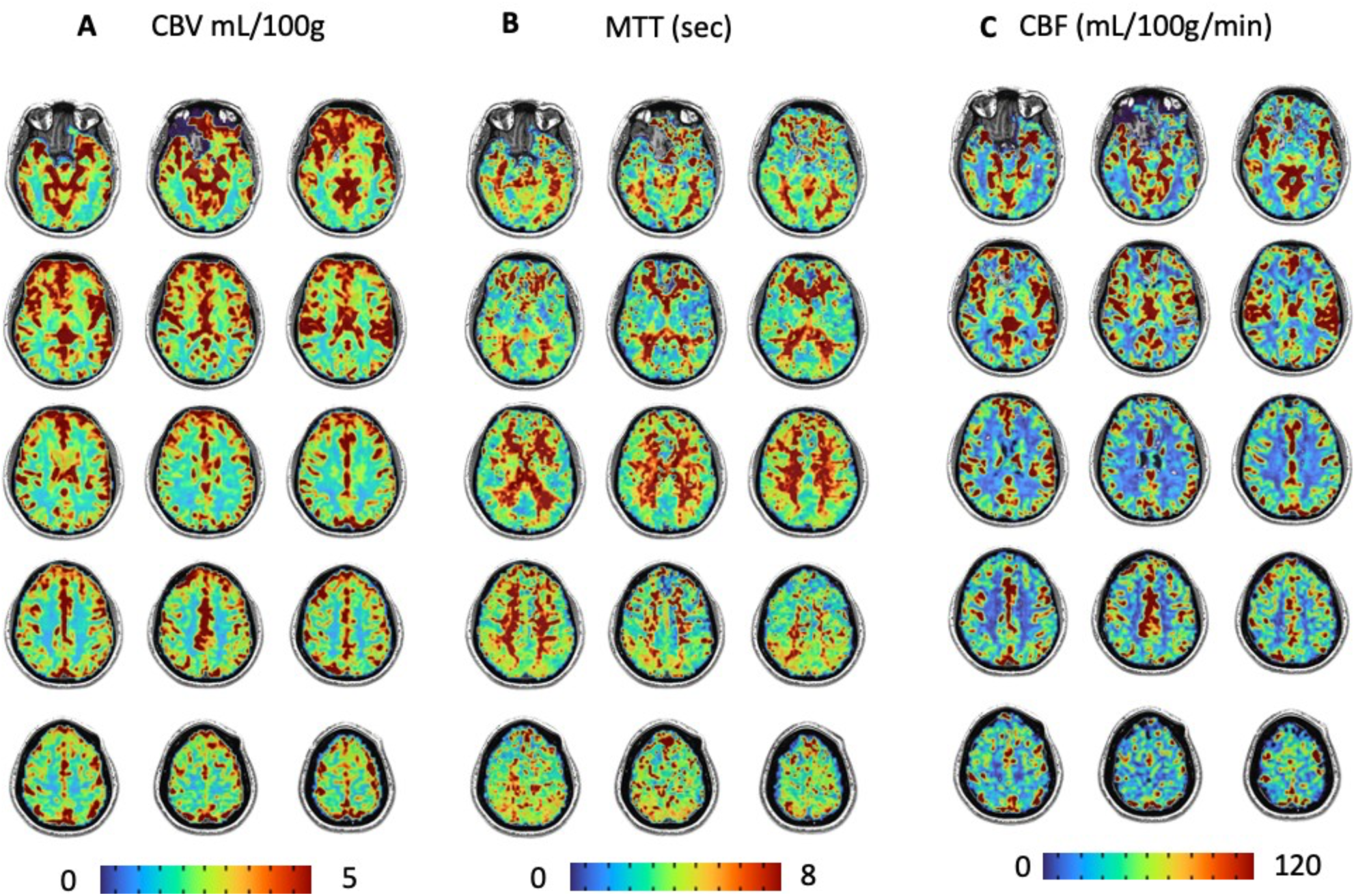
Perfusion metrics Cerebral blood volume (CBV), (mean transit time (MTT), cerebral blood flow (CBF).

## Discussion

We report remarkable findings in the course of protocol development for studies in using [dOHb] as a contrast agent. The main findings are that steady baseline levels of hypoxia and hemoglobin desaturation can be maintained (Figure 1), and that ventilatory challenges can induce an abrupt decrease in [dOHb] in the course of 10s, followed by a rapid return to the baseline saturation (Figure 1). This rapid reduction in [dOHb] is reflected in a rapid increase in BOLD signal with high CNR throughout the arterial and venous compartments of the brain (Figure 2d). The rapidity of the change in BOLD implies a rapid change in [dOHb] in the lung and little dispersion of the bolus prior to arrival in the brain vasculature. The degree of dispersion observed would compare favorably to that of intravenously injected contrast agents as it eliminates an entire pathway over which dispersion can develop including, the brachial vein, the subclavian vein, the right atrium, and the right ventricle. Rapid contrast generation in the lung bypasses this segment of the dispersive pathway, with subsequent dispersion only occurring due to passage within the left atrium and left ventricle with the latter dispersion depending on the left ventricular ejection fraction. The respiratory source of contrast generation therefore enables an arterial contrast bolus as close to a square wave input function as possible short of direct contrast injection into large arteries.

As expected, the rise in the arterial signal precedes the rise in the venous signal. The larger rise in the venous BOLD signal is consistent with a lower baseline of the unscaled BOLD signal. The reason for this is that large veins/sinuses are bigger than large arteries, and more likely to contain an imaging voxel with 100% blood that has high [dOHb] and therefore low signal than voxels over arteries where voxels that include both arterial blood and adjacent tissue that has only 4% blood volume. Therefore when scaling the image, dividing by the lower baseline in veins yields a higher percent change in BOLD signal over veins.

Induction of hyperoxia has been used by others to induce BOLD signal changes to enable measurement of CBF, CBV and MTT^17^. The signal characteristics with hyperoxia differ from intravascular bolus contrast or from abrupt inhalations as in our method.

The hyperoxic inhalation does not change the [dOHb] in the arteries, rather it causes more O_2_ to dissolve into the plasma. On arrival to the tissues, the oxygen dissolved in the plasma is spares that bound to hemoglobin. This mechanism does not generate an arterial input function (AIF). There is however a reduction in [dOHb] in tissue blood, minimally increasing the tissue BOLD signal. Our method changes the [dOHb] in the arterial blood thereby inducing a measurable AIF. As the oxygen content of the blood changes, the effects are passed on to the tissues and the BOLD signal reflects the net [dOHb]. Whereas methods where ventilation and thus PaCO_2_ and thereby cerebral blood flow^18^ may change, with our method cerebral blood flow remains constant over the entire PaO_2_ range of 100 mmHg to 45 mmHg.^19^

Finally, in terms of safety to exposure to brief periods of hypoxia, there is no evidence that it is harmful. For example, in La Paz, the capital city of Bolivia at 4000 m elevation, most of the population, as well as visitors have PO2 in the range of 50-60 mmHg, which may be associated with some limitation in exercise tolerance, but not serious illness. For further perspective, most of the human population undergoes multiple (5-25) episodes of apnea (cessation of breathing) during sleep. PaO2 has frequently been observed to fall to values 30-40 mmHg^6^. Thus, the PaO2 of 50-55 mmHg is well tolerated by humans and frequently encountered as part of day-to-day living. In our study, the low PaO_2_ at baseline was virtually imperceptible, causing no distress to any of the 5 subjects.

In this study, hypoxia was established as a baseline, and a normoxic lung changes with decreasing [dOHb] used as the stimulus. The same imaging principles would apply in the reverse situation with a normoxic baseline and hypoxic gas challenges with increasing [dOHb]. This latter approach would reduce safety concerns in some patients with co-morbidities. The hypoxic baseline was chosen in these studies because the conditions for rapid transition to normoxia were more readily attained at the time of the study than the reverse. Implementation of the study using normoxia as the baseline is planned in continuing studies.

We conclude that the use of [dOHb] changes as a non-invasive intra-vascular MRI contrast agent is feasible.

## Acknowledgement

The study was supported by the Andreae Vascular Dementia Research Unit in the Joint Department of Medical Imaging at The University Health Network and a research grant to Dr. Kamil Uludag from the Institute for Basic Science, Suwon, Republic of Korea (IBS-R015-D1).

